# Comparison of IL-33 and IL-5 Family Mediated Activation of Human Eosinophils

**DOI:** 10.1101/645275

**Authors:** Evelyn L. Angulo, Elizabeth M. McKernan, Paul S. Fichtinger, Sameer K. Mathur

## Abstract

Eosinophils are the prominent inflammatory cell involved in allergic asthma, atopic dermatitis, eosinophilic esophagitis, and hypereosinophilic syndrome and are found in high numbers in local tissue and/or circulating blood of affected patients. There is recent interest in a family of alarmins, including TSLP, IL-25 and IL-33, that are epithelial-derived and released upon stimulation of epithelial cells. Several genome wide association studies have found SNPs in genes encoding IL-33 to be risk factors for asthma. In two studies examining the direct role of IL-33 in eosinophils, there were differences in eosinophil responses. We sought to further characterize activation of eosinophils with IL-33 compared to activation by other cytokines and chemokines. We assessed IL-33 stimulated adhesion, degranulation, chemotaxis and cell surface protein expression in comparison to IL-3, IL-5, and eotaxin-1 on human eosinophils. Our results demonstrate that IL-33 can produce as potent or more eosinophil activation than IL-3, IL-5 and eotaxin-1. Thus, when considering specific cytokine targeting strategies, IL-33 will be important to consider for modulating eosinophil function.

## INTRODUCTION

Eosinophils are the prominent immune cells involved in allergic asthma, atopic dermatitis, eosinophilic esophagitis, and hypereosinophilic syndrome and are found in high numbers in local tissue and/or circulating blood of affected patients (1). In the tissue, eosinophils can release toxic granule contents including Major Basic Protein (MBP), Eosinophil Derived Neurotoxin (EDN), and Eosinophil Peroxidase (EPO), which may cause intended damage to the target in the case of parasitic infections, but can inadvertently damage surrounding host tissue and trigger remodeling. In severe asthma, this can lead to chronic inflammation of the airway resulting in long-term injury and remodeling. In addition, we and others have demonstrated that eosinophils are pro-inflammatory cells signaling other immune cells through cytokine release especially by driving and propagating the Th2 type immune response (2, 3).

IL-5 is a specific activator of eosinophils and is crucial to their development from bone marrow progenitors. It increases adhesion, survival, and cytokine release as well as inhibits apoptosis. However, treatment of asthmatic patients with anti-IL5 drug (Mepolizumab) fails to completely eliminate tissue eosinophilia, though blood eosinophil numbers and asthma exacerbations are significantly reduced (4, 5). Thus, it is important to consider alternative eosinophil activators that may contribute to eosinophil-mediated pathology in airway tissue.

There are other well-studied activators of eosinophils such as IL-3, GM-CSF, and TSLP causing varying degrees of activation and involving different signaling kinetics (6). IL-3 and GM-CSF receptors share the same β chain as IL-5, but IL-3 and GM-CSF have differential effects on eosinophils likely due to the regulation of their specific α chain on eosinophils surface. Eosinophils from bronchial lavage (BAL) have increased expression of IL-3Rα and GM-CSFRα and decreased expression of IL-5Rα compared to circulating blood eosinophils (7). IL-5 added to eosinophils *in vitro* leads to up-regulation of IL-3Rα and GM-CSFRα and down-regulation of IL-5Rα (8, 9). IL-3 more strongly induces eosinophil proteins including CD48, CD13, and Semaphorin 7A than GMCSF and IL-5 (10, 11). When IL-3 is added along with TNFα, mRNA for MMP-9 and Activin A are more strongly increased by transcription and mRNA stability than with IL-5 and GMCSF stimulation (12, 13).

There is more recent interest in a family of alarmins, including TSLP, IL-25 and IL-33. The alarmins are epithelial-derived, and released in response to a variety of triggers including epithelial trauma, allergic inflammation, protease activity, and rhinovirus infection (14, 15). These cytokines are involved in the pathophysiology of allergic diseases, including asthma and atopic dermatitis, through Th2 pathway activation (16, 17). We and others have shown that TSLP can activate eosinophils leading to production of Th2 cytokines, enhanced survival and degranulation (18, 19).

Several genome wide association studies have found SNPs in genes encoding IL-33 or its receptor ST2 (IL1R1) to be risk factors for eosinophilic asthma, early childhood onset asthma, and severe forms of the asthma (20) Airway biopsies of asthmatics have higher IL-33 mRNA expression than healthy subjects(21). In addition, Rhinovirus infection which is an important risk factor for asthma exacerbations, increases IL-33 levels in nasal lavage of asthmatic patients (22). In two studies examining the role of IL-33 in eosinophils, there were differences in eosinophil responses. Cherry et al. demonstrated that IL-33 was as potent or more so than IL-5 in inducing adhesion and degranulation, but did not evaluate chemotaxis (23). Suzukawa et al. demonstrated that IL-33 was more potent than IL-5 in inducing adhesion, but not degranulation (24). In addition, CD11b(αM intergrin) was also shown to have increased expression in the presence of IL-33, but no chemotactic effect was observed by Suzukawa et al. We sought to further characterize activation of eosinophils with IL-33 compared to activation by other cytokines and chemokines.

## METHODS

### Eosinophil purification

In an IRB approved protocol (UW-HS-IRB-2013-1570), adult volunteers (18-55 years) with allergic rhinitis or mild atopic asthma were consented to provide peripheral blood samples (300 ml) in heparinized syringes. Blood was diluted 1:1 with 1x HBSS (Corning), layered over 1.090g/mL Percoll, and spun at 2000rpm for 20 minutes. The mononuclear cell layer and Percoll layers were removed and the red blood cell and granulocyte pellet was moved to a clean tube. Contaminating erythrocytes were removed through two hypotonic lyses with ddH2O and divided into 200 million cell aliquots. The aliquots were incubated at 4°C for 30 minutes with 200µL HBSS with 4% NCS (Sigma), 200µL of anti-CD16 beads, 15µL of anti-CD14 and anti-CD3 beads, and 30µL of anti-glycophorin-A beads. The diluted cells were passed through an Auto MACS (Miltenyi) to remove all of the CD16, CD14, CD3, and glycophorin-A positive cells. Purified cells were 97% – 99.9% eosinophils with neutrophils and lymphocytes as contaminating cells.

### Eosinophil adhesion

Eosinophil adhesion was assessed as previously described by measuring Eosinophil Peroxidase activity (EPO) from adhered cells on 96 well plates (Immulon 4 HBX) stimulated with the respective cytokines for 30 min (25). Wells were coated with 10 ug/ml I-CAM1 (RnD Systems), 5ug/ml V-CAM1(RnD Systems) or 5ug/ml Periostin (RnD Systems). Wells were blocked with 0.1% gelatin in HBSS containing Ca^2^+ and Mg^2^+ and washed with 3 times with HBSS. Cells, 1×10^4^ eosinophils, were added to each well along with the respective cytokines (1ng/ml final concentration). After 30 min at 37°C, wells were washed 3x with HBSS to remove non adherent cells. 100ul of HBSS was added to sample wells and 100ul with 1×10^4^ eosinophils were added to untreated wells to provide “Total” eosinophils for comparison. Eosinophil peroxidase (EPO) activity was used to provide an index of eosinophil cell numbers. EPO substrate (1mN H_2_O_2_, 1mM O-phenylenediamine (Sigma), and 0.1% Triton ×-100 (Sigma) in Tris buffer, pH8.0) was added to each well for 30 minutes at RT before stopping the reaction using 4M H_2_SO_4_. The colorimetric change at 492nm was assessed with the BioTek Synergy HT plate reader and the % adherence was calculated using: sample well OD 492/Total well OD 492 x100%.

### Eosinophil degranulation

Eosinophil degranulation was assessed by performing ELISA for Eosinophil Derived Neurotoxin (EDN) on cell free supernatants from eosinophils cultured for 4hrs with *N*-Formylmethionyl-leucyl-phenylalanine (FMLP) (Calbiochem) or cytokines. Eosinophils in HBSS with 0.03% gelatin and Ca^2^+/Mg^2^+, 2.5 ×10^5^/well, were stimulated with 100nM FMLP as a positive control and IL-5 (RnD Systems), IL-3 (RnD Systems), and IL-33 (RnD Systems) at 1ng/ml final concentration for 4 hours at 37°C. A “Total” EDN content well was created by adding 2.5×10^5^ cells directly to 0.1M HCL + 1% Triton x-100 to lyse the cells and their granules. Cell free supernatants were collected and analyzed with EDN ELISA kit per manufacturer’s instructions (MBL).

### Eosinophil chemotaxis

Eosinophil chemotaxis was measured with a 24-well plate and a 6.5mm thick transwell with 5µm diameter pores. For each condition, 600µL of co-culture media and chemoattractant was added in duplicates to the lower chamber. Eotaxin-1 (Promokine) was used at a concentration of 100ng/mL while IL-3 (RnD Systems), IL-5 (RnD Systems), or IL-33 (RnD Systems) were added at a concentrations of 10ng/mL. The upper transwell was pre-wet using HBSS with 0.01% gelatin and Ca^2^+/Mg^2^+. Purified eosinophils were also diluted in the HBSS with 0.01% gelatin and Ca^2^+/Mg^2^+ to a concentration of 3×10^6/mL. Each upper well of the transwell was aliquoted 100µL of the eosinophil solution and the plate incubated at 37°C for one hour. In order to remove any cells stuck to the bottom of the transwell, 250mM EDTA (Boston Bioproducts) was added to the bottom well for 5 minutes at room temperature. The transwell was then removed and 50µL of the solution from the bottom chamber was then diluted 1:1 with trypsin (0.4%, Gibco) and counted.

### Flow cytometry

Eosinophils purified through AutoMACS separation were incubated with IL-3(10ng/mL, RnD Systems), IL-5(10ng/mL, RnD Systems), IL-33(10ng/mL, RnD Systems), Eotaxin-1(100ng/mL, Promokine), in FBS and 1% RPMI for 4 hours at 37°C (0.5×10^6^ cells/tube). Cells were incubated with antibody-fluorochrome conjugates for 30 minutes at 4°C. Cells were washed, fixed with a paraformaldehyde solution, and analyzed using the BD LSR Fortessa or BD LSR II (BD Biosciences). Antibodies used were: CD16 PE-CF594, CD14 Alexa Fluor 488(BD Biosciences); CD18 PE-Cy7, CD11b PerCP Cy5.5, ICAM-1 PE (Biolegend); CD66b PE-Cy7 (eBioscience); Ghost Dye™ Red 780 (Tonbo). Data was collected using FACSDivaTM Software (BD Biosciences). Data was analyzed with Flow Jo Version 10 (Tree Star Inc.), with 75-150,000 events captured. Experiments were performed with at least 7-9 replicates for each group. Appropriate compensation controls were performed using antibodies and compensation control beads (Invitrogen™ Ebioscience™ UltraComp Ebeads™).

### Statistics

Adhesion, degranulation, and chemotaxis data are expressed as mean ± SEM. One-way ANOVA test and the Dunn’s test or Student Newman Keul’s test, if appropriate were used for pair-wise comparisons (Sigmaplot 13.0, Systat Software). Differences were considered significant when p < 0.05. For flow cytometry studies, MFI values were obtained from flow cytometry analysis software (FlowJo 10.2). MFI fold change values were normalized by performing log_10_ transformation of the ratio between cytokine stimulated MFI values and unstimulated MFI values. Ratio-*t*-tests were performed on the log-transformed data (RStudio Version 1.0.153). One-way ANOVA tests and the Dunn’s test or Student Newman Keul’s test, if appropriate, were used for pair-wise comparisons of the log-transformed data (SigmaPlot 13.0). Differences were considered significant when p < 0.05.

## RESULTS

### Eosinophil Adhesion

Percent adherence was measured by EPO activity from adhered cells (N=22). As shown in Fig 1, eosinophils were significantly more adherent to uncoated wells when stimulated with IL-33 than when stimulated with IL-3 (p < 0.001) or IL-5 (p < 0.001). Similarly, with ICAM-1 as a substrate, eosinophils stimulated with IL-33 were significantly more adherent then those stimulated with IL-3 (p <0.001) or IL-5 (p < 0.001).

**Fig 1.**
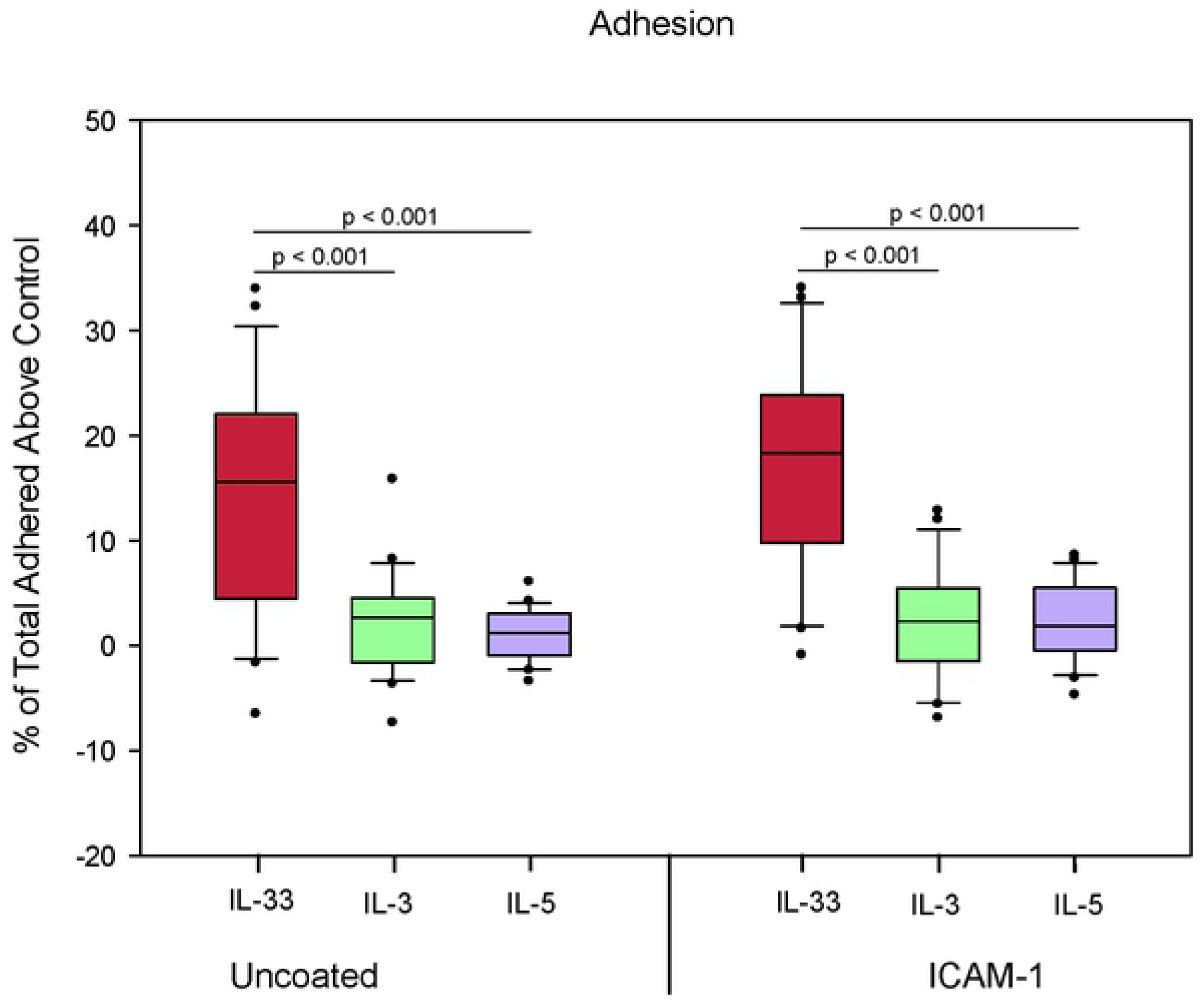
Eosinophils are more adherent to uncoated wells when stimulated with IL-33. Eosinophil adhesion determined by EPO release from adherent cells cultured for 30min with cytokine stimulation, cytokine concentration 1ng/mL. Values shown are a percentage of optical density (OD) of “Total” wells containing 10×10e4 lysed eosinophils with control (no stimulation) subtracted. N=22 Experiments from different blood donors (n=22) with the exception of 3 donors that were repeated twice in which case, results were averaged from the 2 different experiments using their cells. Lines connect comparison groups with p-value denoting significant difference in pair-wise comparisons.

### Eosinophil Degranulation

As shown in Fig 2, eosinophils stimulated with IL-33 for 4 hours induced more percent total EDN release than the positive control FMLP (p = 0.04, N=16). EDN release from IL-33 stimulated eosinophils was significantly higher than eosinophils stimulated with IL-3 (p < 0.001) and IL-5 (p < 0.001). IL-3 and IL-5 both induced significantly higher degranulation than our control (p < 0.001).

**Fig 2.**
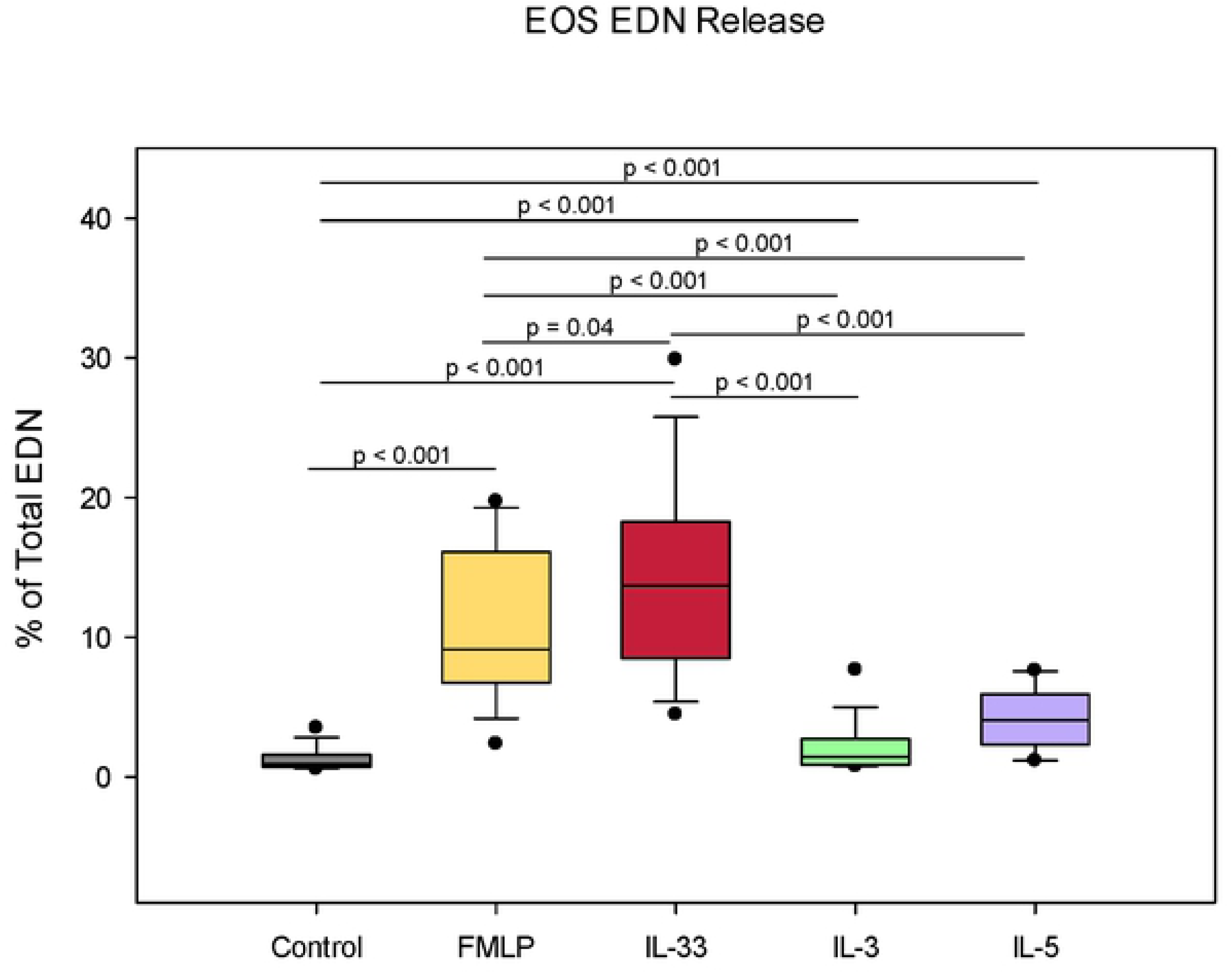
Eosinophils stimulated with IL-33 induce more percent total EDN release than positive control. EDN release determined by ELISA for eosinophils cultured 4 hrs with cytokine stimulation, cytokine concentration 1ng/mL (n=16). Values are expressed as % of “Total” well containing 0.5×10e6/well of lysed eosinophils. Lines connect comparison groups with p-value denoting significant difference in pair-wise comparisons.

### Eosinophil Chemotaxis

As shown in Fig 3, Percent migration of eosinophils after stimulation using transwell chambers was evaluated (N=8). Eotaxin-1 demonstrated significantly higher percent migration than did IL-33 (p < 0.001). IL-3(p < 0.001) and IL-5 (p = 0.005) demonstrated significant increase in percent migration when compared to IL-33. There was no significant difference in percent migration of eosinophils stimulated with IL-33 when compared to control (p = 0.28). Compared to control, percent migration was significantly increased with eotaxin-1 (p < 0.001).

**Fig 3.**
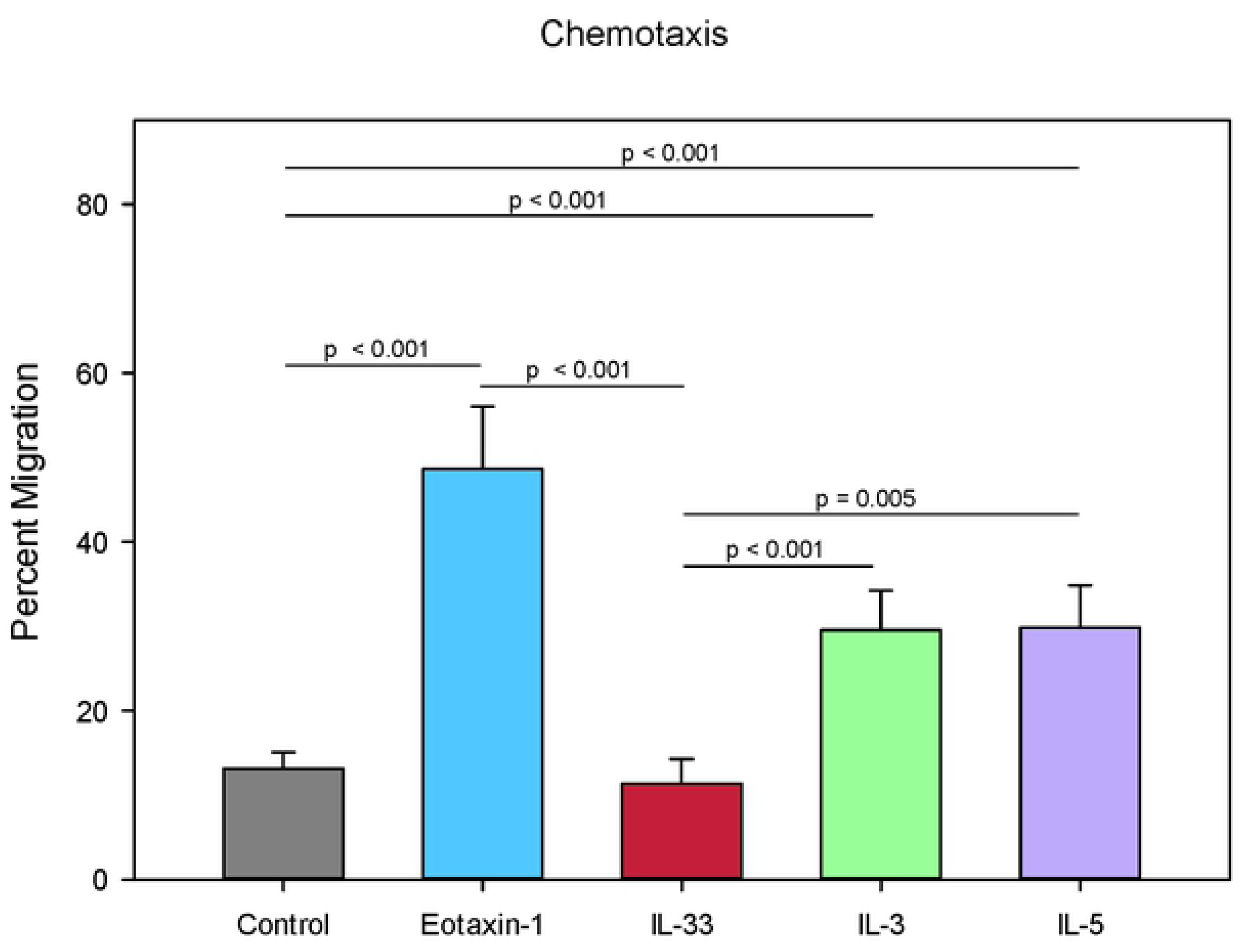
IL-33 stimulation does not result in increased eosinophil migration. Percent migration of eosinophils after stimulation using transwell chambers. Migration assessed after 1 hour after cytokine was placed in lower chamber, eotaxin-1 concentration 100ng/mL, IL-33 and other cytokine concentrations 10ng/mL (n=8). Values are expressed as % migration of total cells in well containing 0.3×10e6/well of eosinophils. Lines connect comparison groups with p-value denoting significant difference in pair-wise comparisons.

### Eosinophil Expression of Cell Surface Markers

As shown in Fig 4, eosinophil cell surface marker expression was assessed using flow cytometry. We compared the change in cytokine stimulated expression for 4 hours versus baseline expression (N=7-9). IL-33, IL-3, IL-5 and eotaxin-1 significantly increased expression of CD11b, CD66b, and CD18 when compared to unstimulated control (with p-values < 0.05). ICAM-1 expression compared to unstimulated control, was significantly increased after incubation with only IL-33, IL-3, IL-5 (with p-values < 0.001), but not eotaxin-1 (p = 0.3). In pairwise comparisons, IL-33 stimulated eosinophils expressed significantly higher levels of CD11b (Fig 4A) and CD18 (Fig 4B) than eotaxin-1 (with p-values < 0.001) or IL-3 stimulated cells (with p-values < 0.05). IL-33 stimulated cells demonstrated higher CD18 expression compared to IL-5 as well (p=0.01). IL-33 stimulated eosinophils significantly increased expression of CD66b (Fig 4C) and ICAM-1 (Fig 4D) when compared to eotaxin-1 (with p-values < 0.001) and IL-3 (with p-values < 0.05), but not IL-5. When comparing IL-33 stimulated eosinophil surface marker expression versus that of IL-3, IL-5 or eotaxin-1 at these concentrations, IL-33 stimulated eosinophils demonstrated the highest fold change in marker expression. The 4hr stimulation experiments did not result in statistically significant change for: N29, CD29, CD11a, CD11c, CD54, CD23, CD40, CD44, CD41, CD62P (Data not shown).

**Fig 4.**
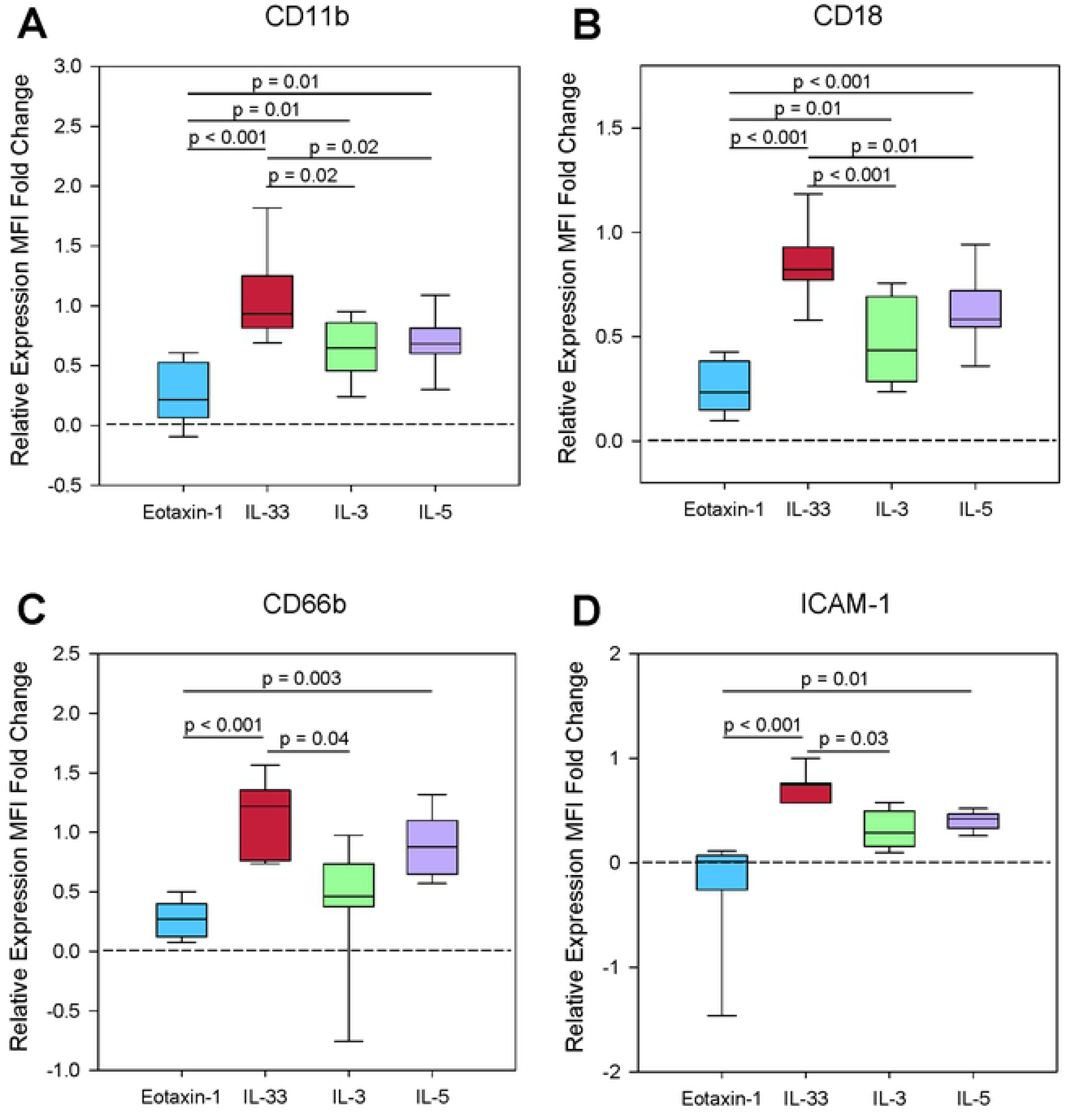
IL-33 induces significant changes in cell surface expression. Box plots of normalized fold change for A) CD11b, B) CD18, C) CD66b, D) ICAM-1 after 4 hour stimulation with either IL-3, IL-5, IL-33, eotaxin-1 compared with unstimulated, represented by dotted line (n=7-8). Lines connect comparison groups with p-value denoting significant difference in pair-wise comparisons.

## DISCUSSION

Our results demonstrate that IL-33 is as or more potent than IL-3 and IL-5 in inducing eosinophil adhesion and degranulation. In addition, eosinophil stimulation with IL-33 resulted in increased expression of the cell surface markers CD11b, CD18, CD66b, and ICAM-1 in a manner comparable to or higher than IL-3, IL-5, and eotaxin-1. Our study confirms the lack of chemotaxis from IL-33 stimulation. Few studies have examined the comparative effects of IL-33 vs. IL-3, IL-5 or eotaxin-1 on eosinophils. To our knowledge, increased eosinophil expression of CD18 and CD66b after IL-33 stimulation has not previously been described.

IL-33 is a member of the IL-1 family of cytokines constitutively expressed in the nucleus of endothelial and epithelial cells of mucosal membranes and fibroblasts and can be released in response to cell injury (26). Characterized as an alarmin, IL-33 warns the immune system of barrier injury when cells undergoing necrosis from infections or physical damage release their contents (27). In this case, the full-length active form is released and IL-33 signaling can occur. However, if cells undergo programmed death like apoptosis, the full length form is cleaved by caspases 3 and 7 and signaling is abrogated (28). Proteases from environmental allergens have also been shown to cleave IL-33, which in turn leads to downstream Th2 signaling and allergic inflammation (29). IL-33 is considered a Th2 type cytokine signaling Th2 lymphocytes and ILC2s to release/produce cytokines such as IL-4, IL-5, IL-6, and IL-13 driving the type 2 immune response. Dendritic cells when stimulated with IL-33, release IL-6 and induce production of IL-5 and IL-13 from naïve CD4+ T cells (30). Viral infections can also result in production of Th2 cytokines. In addition, viral infections can lead to bronchial epithelial damage and trigger the release of inflammatory mediators. In a study by Han et al., mice infected with rhinovirus were found to have increased lung epithelial TSLP and IL-33 (31). The production of these two alarmins, in addition to the presence of Th2 cytokines may result in increased eosinophil activity.

The alarmin family also includes TSLP and IL-25. We have previously demonstrated that TSLP promotes eosinophil degranulation, and that its activity may be enhanced by the allergic cytokine milieu (18). Elevated levels of IL-33 are found in bronchial tissue, and BAL fluid of asthmatics when compared to that of controls (21, 32). This points to an important role for IL-33 in airway inflammation. The IL-33 and TSLP pathways, may be a point of intersection where viral infections and allergic exposures combine to result in increased eosinophilic inflammation and a heightened risk of exacerbation from the activation of eosinophils.

The CD4+ Th2 cytokines IL-3 and IL-5 are produced during allergic inflammation. It has previously been thought that eosinophil activation was mainly controlled by IL-3, IL-5 and GM-CSF. Eosinophil adhesion is the one of the necessary steps in eosinophil migration to target tissues. We observed that eosinophils were significantly more adherent in uncoated or ICAM-1 coated wells after 30 minute incubation with IL-33 when compared to IL-3 and IL-5. This is similar than the findings of Cherry et al. and Suzukawa et al. who both found a significant increase in eosinophil adhesion with IL-33 when compared to IL-5. Our studies used different lengths of time for eosinophil incubation with stimulant than did the previously described studies. In addition, for our adhesion experiments, we used a lower concentration of 1ng/ml for each cytokine, as opposed to the 100ng/mL used in the Cherry et al. study and the 1-100ng/mL used in Suzukawa et al.

In our studies, IL-33 induced degranulation of eosinophils significantly more than IL-3 and IL-5. This differs to the results from Suzukawa et al. who also measured EDN after IL-33 incubation. However, our work supports the finding of Cherry et al. that, IL-33 can induce degranulation to a level of at least that of IL-5. One limitation of our experiments is that they were performed at single time points. Previous work has shown that there can be differences in eosinophil activation at different time points. For example, prolonged activation with IL-3 is more potent than IL-5 and GM-CSF in inducing eosinophil expression of specific activation and adhesion molecules (33). This poses the question of whether IL-33 demonstrates similar potency when compared to IL-3, IL-5, or eotaxin-1 at time points not examined in our study.

Our eosinophil chemotaxis studies did not demonstrate increased cell migration after IL-33 incubation. These results are consistent with that of Suzukawa et al. who reported that IL-33 failed to attract eosinophils in their migration study. As demonstrated in our results, IL-33 stimulation led to increased expression of adhesion and migration molecules CD11b, CD18, CD66b and ICAM-1. ICAM-1 is a well known adhesion molecule that binds to Mac-1 and LFA-1 on the surface of cells or endothelium, and has been shown by Suzukawa et al. to increase after IL-33 incubation. Expression of these cell surface molecules was increased with IL-33 to a similar or higher level than IL-3, IL-5 and eotaxin-1. It is possible that IL-33 participates in priming eosinophils to expresses these markers, but does not directly lead to eosinophil migration. Of note, our flow cytometry results revealed changes in expression that are consistent with prior data. For example, others have demonstrated that eosinophils express increased ICAM-1 after incubation with IL-33 (34, 35). Various studies have demonstrated increased expression of CD11b, CD18, CD66b and ICAM-1 in response to IL-5, IL-3, or GM-CSF, although direct comparisons with IL-33 induced expression were not performed for all of these cell surface markers (33, 36).

CD11b (αM integrin) and CD18 (β2 integrin) form the Mac-1 (macrophage integrin, also known as complement receptor 3, CR3) complex. It has previously been reported that Mac-1 plays a key role in eosinophil degranulation and adhesion (37). IL-33 has been demonstrated to increase CD11b expression to levels comparable to that of IL-5 (24), and we have extended these observations to demonstrate that the CD18 dimer partner is also upregulated. CD66b (CEACAM8) is GPI anchored glycoprotein, which has been associated with CD11b. Yoon et al. demonstrated that CD66b cross linking by monoclonal antibody, or galectin-3 led to CD11b clustering on the eosinophil cell surface, and promoted degranulation (36). Therefore, the IL-33 mediated increased CD66b and CD11b likely play a role in enhanced eosinophil degranulation.

These eosinophil cell surface molecules have been studied in a number of inflammatory diseases. Sputum eosinophils of asthma patients have been shown to have up-regulated CD66b and CD11b (38). A study of peripheral eosinophils of asthma patients in response to segmental antigen challenge demonstrated increases in the CD11b and CD18 post-challenge (39). CD66b has been found to be elevated on the surface of peripheral eosinophils in untreated eosinophilic esophagitis patients when compared to healthy controls. Other inflammatory disorders such as rheumatoid arthritis have demonstrated that CD66b, CD11b, and CD18 are down-regulated by glucocorticoid use. These studies highlight that these cell surface markers likely play an important role in mediating inflammation. Our results indicate that IL-33 increased the expression of CD66b, CD18, and CD11b to the level of or more so than IL-3, IL-5 and eotaxin-1. A representative sample of CD11b^high^CD18^high^ and CD11b^high^CD66b^high^ percent positive eosinophils from our stimulation experiments is shown in Table 1. IL-33 induced as high or higher percent-positive cells exhibiting these markers. Therefore, it is plausible that IL-33 leads to potent eosinophil activation through its effects on CD66b, and thereby the Mac-1 complex.

**Table 1.**
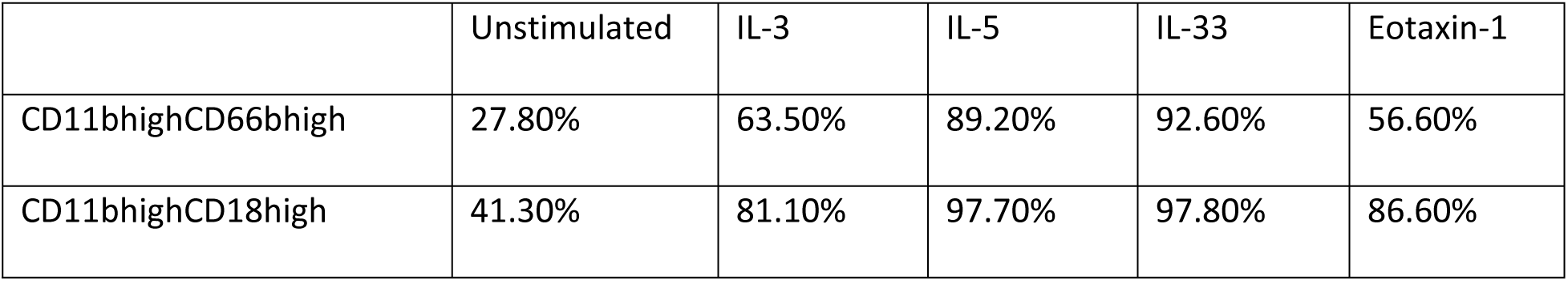
Representative Sample of Double “High” Percent Positive Eosinophils

The above table contains double “high” population percentages for the markers listed. These percentages were obtained by gating on the unstimulated median fluorescence intensity and observing the increase in double “high” percentage shown in a single representative sample.

We recently described relative abundance of proteins of the peripheral blood eosinophil proteome (40). In comparing protein content of IL-5, IL-3 and IL-33 receptor molecules, IL-5Rα was the highest, with the relative abundance of IL-3Rα and ST2 being 15.6% and 28.1% respectively. This demonstrates that despite less relative abundance of the ST2 receptor when compared to IL-5Rα, IL-33 is still capable of inducing comparable or higher eosinophil adhesion, degranulation and cell surface markers when compared to IL-5.

In this paper, we identify that IL-33 exerts its effects on eosinophils, resulting in as potent or more adhesion, degranulation, and cell surface marker expression when compared to IL-3, IL-5, or eotaxin-1. IL-33 as an alarmin has the ability to set off a number of down-stream effects that lead to a skewing toward the Th2 pathway and the development of allergic inflammation. The fact that IL-33 has as potent effects on eosinophils strengthens the argument for use of this molecule as a therapeutic target and broadens our knowledge in regards to how these pathways may be intersecting to produce inflammation. Anti-IL-33 biologics are currently in clinical trials for asthma and other atopic conditions and will be expected to significantly impact eosinophil function in allergic disorders.

## ACKNOWLEDGEMENTS

We thank the nurse coordinators for recruiting patients and providing samples for this study. We also thank the patients who volunteered their time and participated in our study.

## REFERENCES

1. Bochner BS. The eosinophil: For better or worse, in sickness and in health. Annals of allergy, asthma & immunology : official publication of the American College of Allergy, Asthma, & Immunology. 2018;121(2):150–5.

2. Jacobsen EA, Taranova AG, Lee NA, Lee JJ. Eosinophils: Singularly destructive effector cells or purveyors of immunoregulation? Journal of Allergy and Clinical Immunology. 2007;119(6):1313–20.

3. Liu LY, Mathur SK, Sedgwick JB, Jarjour NN, Busse WW, Kelly EAB. Human airway and peripheral blood eosinophils enhance Th1 and Th2 cytokine secretion. Allergy. 2006;61:589–97.

4. Cabon Y, Molinari N, Marin G, Vachier I, Gamez AS, Chanez P, et al. Comparison of antiinterleukin-5 therapies in patients with severe asthma: global and indirect meta-analyses of randomized placebo-controlled trials. Clinical and experimental allergy : journal of the British Society for Allergy and Clinical Immunology. 2017;47(1):129–38.

5. Kelly EA, Esnault S, Liu LY, Evans MD, Johansson MW, Mathur S, et al. Mepolizumab Attenuates Airway Eosinophil Numbers, but Not Their Functional Phenotype in Asthma. Am J Respir Crit Care Med. 2017.

6. McBrien CN, Menzies-Gow A. The Biology of Eosinophils and Their Role in Asthma. Frontiers in medicine. 2017;4:93.

7. Bates ME, Liu LY, Esnault S, Stout BA, Fonkem E, Kung V, et al. Expression of interleukin-5-and granulocyte macrophage-colony-stimulating factor-responsive genes in blood and airway eosinophils. American Journal of Respiratory Cell and Molecular Biology. 2004;30(5):736–43.

8. Liu LY, Sedgwick JB, Bates ME, Vrtis RF, Gern JE, Kita H, et al. Decreased expression of membrane IL-5 receptor alpha on human eosinophils: II. IL-5 down-modulates its receptor via a proteinase-mediated process. Journal of Immunology. 2002;169(11):6459–66.

9. Gregory B, Kirchem A, Phipps S, Gevaert P, Pridgeon C, Rankin SM, et al. Differential regulation of human eosinophil IL-3, IL-5, and GM-CSF receptor alpha-chain expression by cytokines: IL-3, IL-5, and GM-CSF down-regulate IL-5 receptor alpha expression with loss of IL-5 responsiveness, but up-regulate IL-3 receptor alpha expression. J Immunol. 2003;170(11):5359–66.

10. Esnault S, Johansson MW, Kelly EA, Koenderman L, Mosher DF, Jarjour NN. IL-3 up-regulates and activates human eosinophil CD32 and alphaMbeta2 integrin causing degranulation. Clinical and experimental allergy : journal of the British Society for Allergy and Clinical Immunology. 2017;47(4):488–98.

11. Esnault S, Kelly EA, Johansson MW, Liu LY, Han ST, Akhtar M, et al. Semaphorin 7A is expressed on airway eosinophils and upregulated by IL-5 family cytokines. Clinical immunology (Orlando, Fla). 2014;150(1):90–100.

12. Kelly EA, Esnault S, Johnson SH, Liu LY, Malter JS, Burnham ME, et al. Human eosinophil activin A synthesis and mRNA stabilization are induced by the combination of IL-3 plus TNF. Immunology and cell biology. 2016;94(7):701–8.

13. Kelly EA, Liu LY, Esnault S, Quinchia Johnson BH, Jarjour NN. Potent synergistic effect of IL-3 and TNF on matrix metalloproteinase 9 generation by human eosinophils. Cytokine. 2012;58(2):199–206.

14. Han H, Roan F, Ziegler SF. The atopic march: current insights into skin barrier dysfunction and epithelial cell-derived cytokines. Immunol Rev. 2017;278(1):116–30.

15. Mitchell PD, O’Byrne PM. Epithelial-Derived Cytokines in Asthma. Chest. 2017;151(6):1338–44.

16. Al-Sajee D, Oliveria JP, Sehmi R, Gauvreau GM. Antialarmins for treatment of asthma: future perspectives. Current opinion in pulmonary medicine. 2018;24(1):32–41.

17. Ziegler SF, Artis D. Sensing the outside world: TSLP regulates barrier immunity. Nat Immunol. 2010;11(4):289–93.

18. Cook EB, Stahl JL, Schwantes EA, Koziol-White CJ, Bertics PJ, Graziano FM, et al. Thymic Stromal Lymphopoietin Directly Activates Eosinophil Degranulation. The Journal of Allergy and Clinical Immunology. 2010;125(2):AB133.

19. Wong CK, Hu S, Cheung PFY, Lam CW. TSLP Induces Chemotactic and Pro-survival Effects in Eosinophils: Implications in Allergic Inflammation. American Journal of Respiratory Cell and Molecular Biology. 2010;43(3):305–15.

20. Jones BL, Rosenwasser LJ. Linkage and Genetic Association in Severe Asthma. Immunol Allergy Clin North Am. 2016;36(3):439–47.

21. Prefontaine D, Nadigel J, Chouiali F, Audusseau S, Semlali A, Chakir J, et al. Increased IL-33 expression by epithelial cells in bronchial asthma. J Allergy Clin Immunol. 2010;125(3):752–4.

22. Jackson DJ, Makrinioti H, Rana BM, Shamji BW, Trujillo-Torralbo MB, Footitt J, et al. IL-33-dependent type 2 inflammation during rhinovirus-induced asthma exacerbations in vivo. Am J Respir Crit Care Med. 2014;190(12):1373–82.

23. Cherry WB, Yoon J, Bartemes KR, Iijima K, Kita H. A novel IL-1 family cytokine, IL-33, potently activates human eosinophils. J Allergy Clin Immunol. 2008;121(6):1484–90.

24. Suzukawa M, Koketsu R, Iikura M, Nakae S, Matsumoto K, Nagase H, et al. Interleukin-33 enhances adhesion, CD11b expression and survival in human eosinophils. Laboratory investigation; a journal of technical methods and pathology. 2008;88(11):1245–53.

25. Sedgwick JB, Calhoun WJ, Vrtis RF, Bates ME, Mcallister PK, Busse WW. Comparison of Airway and Blood Eosinophil Function After Invivo Antigen Challenge. Journal of Immunology. 1992;149(11):3710–8.

26. Moussion C, Ortega N, Girard JP. The IL-1-like cytokine IL-33 is constitutively expressed in the nucleus of endothelial cells and epithelial cells in vivo: a novel ‘alarmin’? PLoS One. 2008;3(10):e3331.

27. Mehraj V, Ponte R, Routy JP. The Dynamic Role of the IL-33/ST2 Axis in Chronic Viral-infections: Alarming and Adjuvanting the Immune Response. EBioMedicine. 2016;9:37–44.

28. Luthi AU, Cullen SP, McNeela EA, Duriez PJ, Afonina IS, Sheridan C, et al. Suppression of interleukin-33 bioactivity through proteolysis by apoptotic caspases. Immunity. 2009;31(1):84–98.

29. Cayrol C, Duval A, Schmitt P, Roga S, Camus M, Stella A, et al. Environmental allergens induce allergic inflammation through proteolytic maturation of IL-33. Nat Immunol. 2018;19(4):375–85.

30. Rank MA, Kobayashi T, Kozaki H, Bartemes KR, Squillace DL, Kita H. IL-33-activated dendritic cells induce an atypical TH2-type response. J Allergy Clin Immunol. 2009;123(5):1047–54.

31. Han M, Rajput C, Hong JY, Lei J, Hinde JL, Wu Q, et al. The Innate Cytokines IL-25, IL-33, and TSLP Cooperate in the Induction of Type 2 Innate Lymphoid Cell Expansion and Mucous Metaplasia in Rhinovirus-Infected Immature Mice. J Immunol. 2017;199(4):1308–18.

32. Li Y, Wang W, Lv Z, Chen Y, Huang K, Corrigan CJ, et al. Elevated Expression of IL-33 and TSLP in the Airways of Human Asthmatics In Vivo: A Potential Biomarker of Severe Refractory Disease. J Immunol. 2018;200(7):2253–62.

33. Esnault S, Kelly EA. Essential Mechanisms of Differential Activation of Eosinophils by IL-3 Compared to GM-CSF and IL-5. Crit Rev Immunol. 2016;36(5):429–44.

34. Chow JYS, Wong CK, Cheung PFY, Lam CWK. Intracellular signaling mechanisms regulating the activation of human eosinophils by the novel Th2 cytokine IL-33: implications for allergic inflammation. Cellular And Molecular Immunology. 2009;7:26.

35. Wong C-K, Leung KM-L, Qiu H-N, Chow JY-S, Choi AOK, Lam CW-K. Activation of Eosinophils Interacting with Dermal Fibroblasts by Pruritogenic Cytokine IL-31 and Alarmin IL-33: Implications in Atopic Dermatitis. PLOS ONE. 2012;7(1):e29815.

36. Yoon J, Terada A, Kita H. CD66b regulates adhesion and activation of human eosinophils. J Immunol. 2007;179(12):8454–62.

37. Horie S, Kita H. CD11b/CD18 (Mac-1) is required for degranulation of human eosinophils induced by human recombinant granulocyte-macrophage colony-stimulating factor and platelet-activating factor. J Immunol. 1994;152(11):5457–67.

38. Tak T, Hilvering B, Tesselaar K, Koenderman L. Similar activation state of neutrophils in sputum of asthma patients irrespective of sputum eosinophilia. Clin Exp Immunol. 2015;182(2):204–12.

39. Johansson MW, Gunderson KA, Kelly EA, Denlinger LC, Jarjour NN, Mosher DF. Anti-IL-5 attenuates activation and surface density of β(2) -integrins on circulating eosinophils after segmental antigen challenge. Clin Exp Allergy. 2013;43(3):292–303.

40. Wilkerson EM, Johansson MW, Hebert AS, Westphall MS, Mathur SK, Jarjour NN, et al. The Peripheral Blood Eosinophil Proteome. Journal of Proteome Research. 2016;15(5):1524–33.

